# Natural statistics of head roll: implications for Bayesian inference in spatial orientation

**DOI:** 10.1101/2022.09.07.506906

**Authors:** Sophie C.M.J. Willemsen, Leonie Oostwoud Wijdenes, Robert J. van Beers, Mathieu Koppen, W. Pieter Medendorp

**Author notes:** **Corresponding author:** Sophie Willemsen, Radboud University Nijmegen, Donders Institute for Brain, Cognition and Behaviour, Thomas van Aquinostraat 4, 6525 GD Nijmegen, The Netherlands.

## Abstract

We previously proposed a Bayesian model of multisensory integration in spatial orientation (1). Using a Gaussian prior, centered on an upright head orientation, this model could explain various perceptual observations in roll-tilted participants, such as the subjective visual vertical, the subjective body tilt (1), the rod-and-frame effect (2), as well as their clinical (3) and age-related deficits (4). Because it is generally assumed that the prior reflects an accumulated history of previous head orientations, and recent work on natural head motion suggests non-Gaussian statistics, we examined how the model would perform with a non-Gaussian prior. In the present study, we first experimentally generalized the previous observations in showing that also the natural statistics of head orientation are characterized by long tails, best quantified as a *t*-location-scale distribution. Next, we compared the performance of the Bayesian model and various model variants using such a *t*-distributed prior to the original model with the Gaussian prior on their accounts of previously published data of the subjective visual vertical and subjective body tilt tasks. All of these variants performed substantially worse than the original model, suggesting a special value of the Gaussian prior. We provide computational and neurophysiological reasons for the implementation of such a prior, in terms of its associated precision–accuracy trade-off in vertical perception across the tilt range.

**New & Noteworthy:** It has been argued that the brain uses Bayesian computations to process multiple sensory cues in vertical perception, including a prior centered on upright head orientation which is usually taken to be Gaussian. Here, we show that non-Gaussian prior distributions, although more akin to the statistics of head orientation during natural activities, provide a much worse explanation of such perceptual observations than a Gaussian prior.

## Introduction

Sensory systems are thought to be optimized for processing naturalistic stimuli (5–8). Given uncertainty in the moment-to-moment sensory information, the statistical regularities within the sensory environment, which can be inferred from an accumulated history of the system’s previous sensory states, add informational value to creating perception. For example, it has been shown that the ‘light-comes-from-above’ experience is used to interpret complex and ambiguous visual input (9) and that the predominance of horizontal and vertical orientations in natural scenes is used in visual orientation perception (10).

Bayesian theory provides a formal framework to describe sensory processing under uncertainty. According to this theory, next to the available sensory evidence also a default assumption about the state, expressed in the form of a prior distribution, is taken into account. Bayes’ rule is the statistically optimal way to combine this prior with noisy sensory information. In lab-based paradigms, the prior often accounts for otherwise unexplainable biases (11, 12). While the prior distribution can be of any type (10, 13), it is often assumed to be a Gaussian distribution for reasons of computational convenience (14, 15).

Earlier work from our lab has proposed a Bayesian model of multisensory integration for spatial orientation (1). In this model we assumed that, to process vestibular and other sensory information, the brain uses a Gaussian prior centered on upright. Based on this prior, the model could explain the well-known Aubert effect – the underestimation of head tilt – when the head is roll-oriented using a vestibular chair (16–19). In subsequent studies, we showed that this model could also explain age-related sensory reweighting in spatial orientation (4), certain behavioral observations in patients (3), and visual contextual effects on spatial orientation (2). The model could also explain vertical perception in monkeys and proprioceptive reweighting following complete vestibular loss (20). However, whether the Gaussian prior in this model reflects the statistics of head orientation during natural activities is unclear.

There are studies that suggest that natural motion statistics are typically described by non-Gaussian distributions (6, 21, 22). For example, Carriot et al. (6) recorded the head’s angular velocity and linear acceleration while participants performed everyday movements such as walking, running, or riding a bus. Measured probability distributions of the head’s angular velocity and linear acceleration were not Gaussian but had long tails as quantified by large positive excess kurtosis values. Hausamann et al. (21) measured head and trunk movements for long durations (>10 h) without explicit instructions and reported skewed acceleration distributions.

Building further on this work, in the present study we test the hypothesis that the Aubert effect in spatial orientation is explained by a prior that corresponds to the statistics of head orientation during natural activities. Adding to and generalizing the existing literature about the statistics of natural head motion, we first recorded head orientation in human participants while they performed everyday movements, and calculated probability density distributions of head orientation in space. The kurtosis values obtained generally indicated clearly non-Gaussian distributions. Next, the original Gaussian model by Clemens et al. (1) with a closed-form solution was converted into a numerical model to enable computations with non-Gaussian priors. This numerical model was fit to the previously obtained psychometric data on spatial orientation to test whether alternative real-world priors account for lab-derived Aubert effects. Since all data are interpreted within the general structure of the model by Clemens et al. (1), we begin with a short modeling background.

### Modeling background

Clemens et al. (1) proposed a Bayesian model of the transformation and integration of various sensory signals (from body, head, and neck) into two spatial orientation estimates: the subjective body tilt (SBT) and subjective visual vertical (SVV) (see Figure 1). The sensory signals considered are body orientation in space from tactile receptors in the skin, head orientation in space as be provided by the otoliths, and head orientation relative to the body by neck proprioception. These sensory signals are represented by Gaussian distributions. The neck signal provides a transformation between body orientation and head orientation, thus creating indirect sources of information for both estimates. Final optimal estimates involve the integration of direct and indirect information as well as prior information.

**Figure 1.**
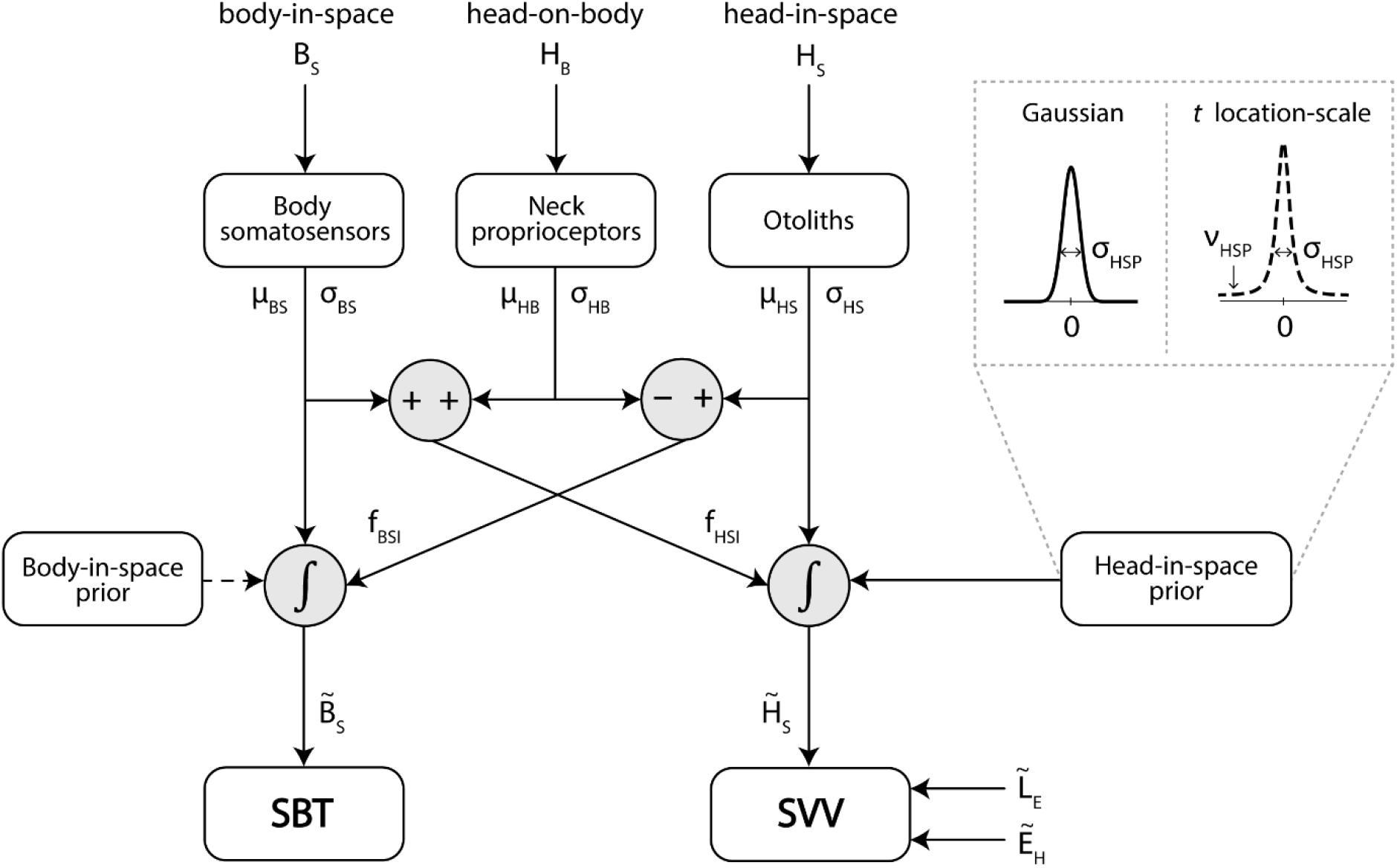
Schematic representation of the sensory integration model by Clemens et al. (1). Body somatosensors, neck proprioceptors and otoliths measure the orientation of the body in space (*B_S_*), the head on the body (*H_B_*) and the head in space (*H_S_*), respectively. The neck signal enables a reference frame transformation of the body-tilt signal into a head-in-space signal (*f_HSI_*) and a transformation of the head-tilt signal into a body-in-space signal (*f_BSI_*). To compute the optimal estimate of body orientation 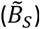, the body-tilt signal is combined with the transformed head-tilt signal, assuming a uniform body-in-space prior. The optimal estimate of head-in-space orientation 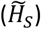 is determined by integration of the otolith signal, the transformed body-in-space signal and a head-in-space prior. The original model by Clemens et al. (1) represents the head-in-space prior as a Gaussian distribution with a mean fixed at 0 and standard deviation *σ_HSP_*. In the current study, we consider a *t*-location-scale distribution with a location parameter fixed at 0, scale parameter *σ_HSP_* and shape parameter *v_HSP_* as head-in-space prior. To acquire an estimate of the line-in-space orientation, 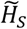 is combined with estimates of the eye-in-head 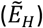 and line-on-eye 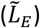 orientation.

Clemens et al. (1) took all three sensory distributions to be unbiased Gaussians, i.e., with a mean equal to the actual tilt angle. The standard deviations of the body sensor and neck sensor constitute two free parameters *σ_BS_* and *σ_HB_*, respectively, while the standard deviation of the otolith signal is assumed to increase linearly with absolute tilt angle (12, 23), requiring another two free parameters *a_HS_* and *b_HS_*, with *σ_HS_* = *a_HS_* · |tilt| + *b_HS_*.

Because the SBT data was virtually unbiased across the tilt range, an uninformative, flat body-in-space prior was used, but Clemens et al. (1) included a head-in-space prior centered on zero to account for the systematic underestimation at large tilt angles as observed in the SVV data – the Aubert effect. This prior distribution was also assumed to be Gaussian, with standard deviation *σ_HSP_* as another free parameter.

Since the SVV pertains to the perceived orientation of a visual line, the final head-in-space estimate is to be supplemented with an eye-in-head estimate (*E_H_*), involving the amplitude of the uncompensated ocular counterroll (*A_OCR_*) as a free parameter, and a retinal line orientation estimate (*L_E_*), assumed to be accurate. Both *E_H_* and *L_E_* have small noise levels (<1°), which were ignored. In addition to the six parameters mentioned, the model has a seventh parameter *λ* to account for lapses with an upper bound of 0.06.

In the current study, we focused on the assumption that prior knowledge about head orientation is represented as a Gaussian distribution, and we wanted to test whether the Clemens et al. (1) model can better explain the data using a prior distribution corresponding more closely to the statistics of head orientation in everyday life. Based on previous results (6, 21, 22) as well as newly recorded data (see Methods and Results), we considered the *t*-location-scale distribution as an alternative distribution for the head-in-space prior. The *t*-location-scale distribution is symmetric and unimodal (bell-shaped), like the Gaussian, but it has heavier tails. It has one more parameter than the Gaussian distribution, which influences the shape of the distribution. The *t*-location-scale probability density function is given by

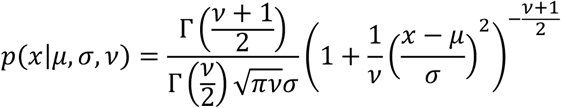

where Γ(·) is the gamma function, *μ* the location parameter, *σ* the scale parameter and *v* the shape parameter (*v* > 0). Smaller values of *v* yield heavier tails; as *v* increases towards infinity, the *t*-location-scale distribution approaches the Gaussian distribution. A direct consequence of this adaptation to the Clemens et al. (1) model is that closed-form expressions for the posterior distribution in terms of the likelihood and prior distributions no longer exist. All computations were therefore done numerically.

## Methods

### Data sets

Two data sets were used to test the model predictions. The first data set, collected previously, is extensively described in Clemens et al. (1). This data collection tabulates psychometric data of seven participants (6 male, 1 female), aged 23-65 years, each performing the SBT and SVV task at different tilt angles, passively imposed by a vestibular chair. Each participant performed 20 experimental sessions of 45 minutes each, yielding over 15 hours of recording time. In short, in the SBT task, participants were first rotated to a randomly chosen tilt angle and then asked to indicate whether their body orientation was CW or CCW from an instructed reference orientation (i.e., either upright (0°), 45° or 90° right side down, or 45° or 90° left side down). Responses were collected using the method of constant stimuli, yielding 140 data points for each instructed reference orientation. The SVV was tested at nine roll-tilt angles, ranging from −120 to 120° at 30° intervals. At each tilt angle, a luminous line was briefly flashed, and the participant indicated whether its orientation in space was CW or CCW from the perceived direction of gravity. The line orientation was selected randomly from a set of 11 line orientations. Each set was tested 12 times, thus yielding a total of 132 data points for each tilt angle. The original model by Clemens et al. (1), which assumed a Gaussian head-in-space prior, provided a very good account of these data.

The second data set was collected anew as a supplement to the existing literature about the statistics of natural head motion (6, 21, 24). Six participants (3 male, 3 female) aged 23-28 years, free of any known neurological or movement disorders, gave written informed consent to track their unconstrained naturalistic motion using inertial measurement units (Xsens MTw Awinda), placed on the pelvis, shoulders, sternum, upper arms, forearms, hands and head. The system was calibrated while the participant was standing in a relaxed, upright position, with their feet parallel to each other and their arms flat against their body, while looking straight ahead with a natural head position. Analogous to Carriot et al. (6), participants performed 5 different naturalistic tasks in and around our university, each one to three times, for 2 min each: walking, running, going up and down the stairs, sitting, and standing. This study was approved by the ethics committee of the Faculty of Social Sciences of Radboud University Nijmegen, the Netherlands. To bring this data set to bear on the Clemens et al. (1) model, we analyzed the roll-tilt angles of the head in space, in degrees. The pre-processing of the raw head orientation data (which was measured in quaternion form) consisted of transforming the raw data to Euler roll-tilt angles in degrees. Head orientation distributions are described in terms of four statistical moments: mean, standard deviation, skewness and kurtosis. To determine which probability distribution best captured the natural head statistics, theoretical distributions were fitted to the head orientation data using Matlab’s built-in Maximum Likelihood Estimation function. We considered the following probability distributions: the Gaussian, logistic, *t*-location-scale (25), extreme value (26) and generalized extreme value distributions (27). The resulting fits were ranked according to their AIC scores.

### Modeling

#### Model implementation

The model was implemented in Matlab (R2019a) and numerically simulated under different assumptions of the prior distribution. Incorporating a non-Gaussian prior into the model caused the closed-form expressions in the original model to no longer exist. Therefore, the probability distributions in the new model implementation were numerically approximated in a circular plane with a resolution of 0.1 degree. A smaller step size did not impact the model predictions but increased the duration of the fitting procedure considerably.

The model was evaluated 500 times for each tilt angle tested in the SBT and SVV tasks. On each of these 500 replications, the mean of each sensory signal (*μ_BS_, μ_HB_* and *μ_HS_*) was randomly drawn from a circular normal distribution, using the actual tilt angle on that trial (*B_S_, H_B_* or *H_S_*) as mean and the standard deviation of the sensory signal (*σ_BS_, σ_HB_* or *σ_HS_*) as standard deviation. The distributions of the indirect signals (*f_BSI_* and *f_HSI_*) were computed following the expressions in Clemens et al. (1) (Equation 2, 4, 6 and 8) as circular normal distributions with means *μ_HS_* – *μ_HB_* and *μ_BS_* + *μ_HB_* and standard deviations 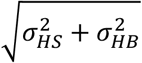 and 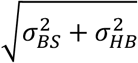, respectively. The body-in-space posterior distribution consists of an integration of the directly and indirectly measured body-in-space information. Similarly, the head-in-space posterior distribution was computed by integrating the (direct and indirect) sensory information and the prior distribution. Taking the mode of the posterior distributions then resulted in the final, optimal estimates (denoted by 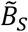 and 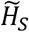 in Figure 1). Note that with the prior in the form of a *t*-location-scale distribution, the resulting posterior is no longer symmetric. Finally, we averaged the modes across the 500 model simulations to determine 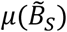 and 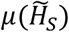 for each tilt angle tested in the two tasks. Similarly, the variance of the modes represents 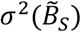 and 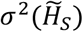. We verified our numerical implementation by fitting the model with a Gaussian prior and found similar model likelihoods and fitted parameter values as via the original estimation procedure used in Clemens et al. (1), validating the new implementation.

#### Model variants and their evaluation

The numerical model version allows to test the model architecture under various assumptions. Models were fitted to the data set from Clemens et al. (1) by minimizing the negative loglikelihood function using the Matlab function ‘fmincon’ (see Clemens et al. (1) for a detailed description of the fitting procedure). We tested various variants of the model with a Gaussian prior (GP models) or a *t*-location-scale prior (TP models), which are summarized in Figure 4. We computed AIC scores to evaluate their performance by accounting for different numbers of free parameters. A lower AIC score indicates a better description of the data by a model variant.

**Figure 2.**
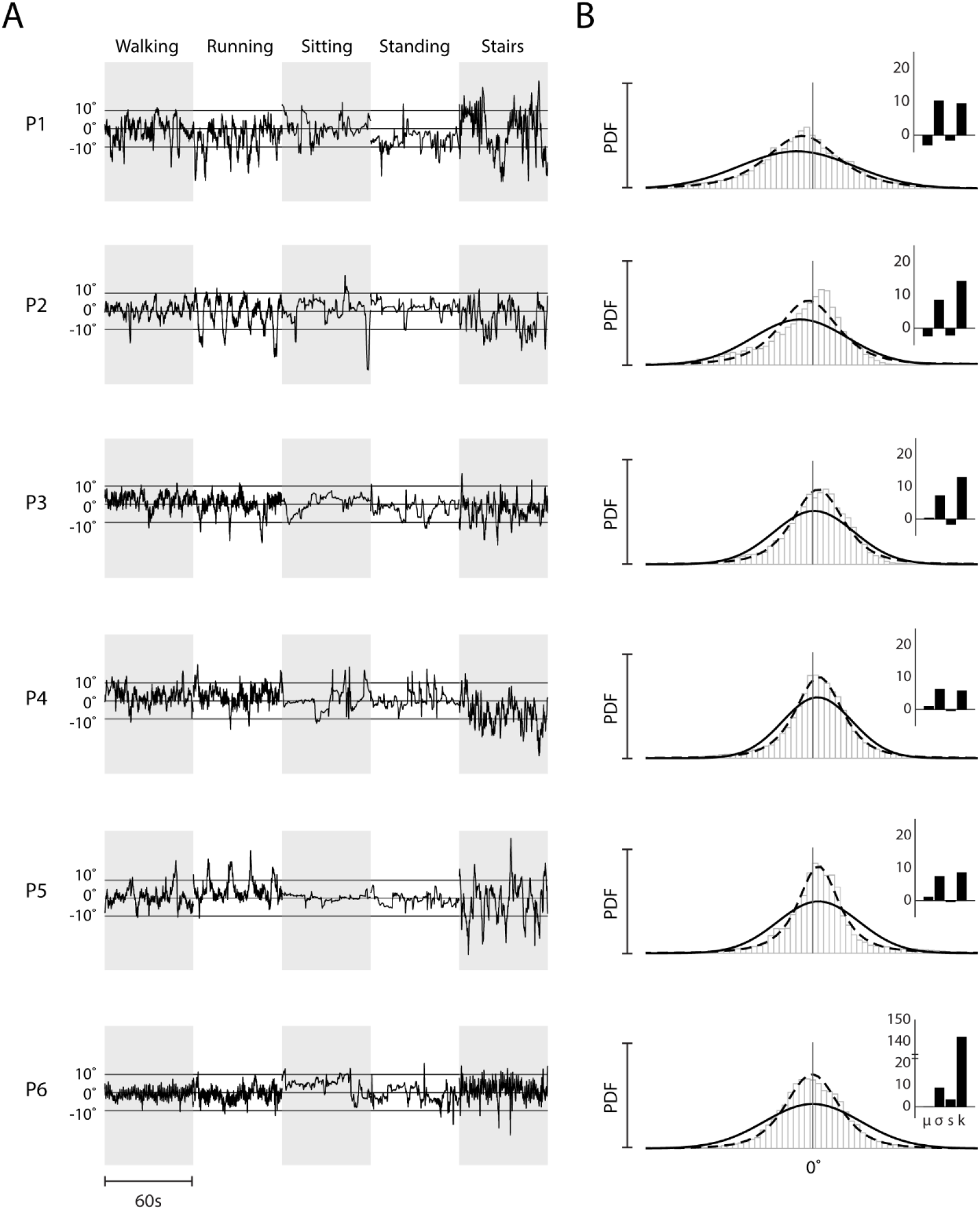
***A***: Representative 60-s traces of the measured head orientations during the different activities, for each participant. ***B***: Fitted normal (solid line) and *t*-location-scale (dashed line) probability density distributions (PDFs), plotted on top of all head roll-tilt data, pooled across activities for each participant in the naturalistic motion tracking experiment. Insets show the four statistical moments (mean *μ*, standard deviation *σ*, skewness *s* and kurtosis *k*) of the pooled data.

**Figure 3.**
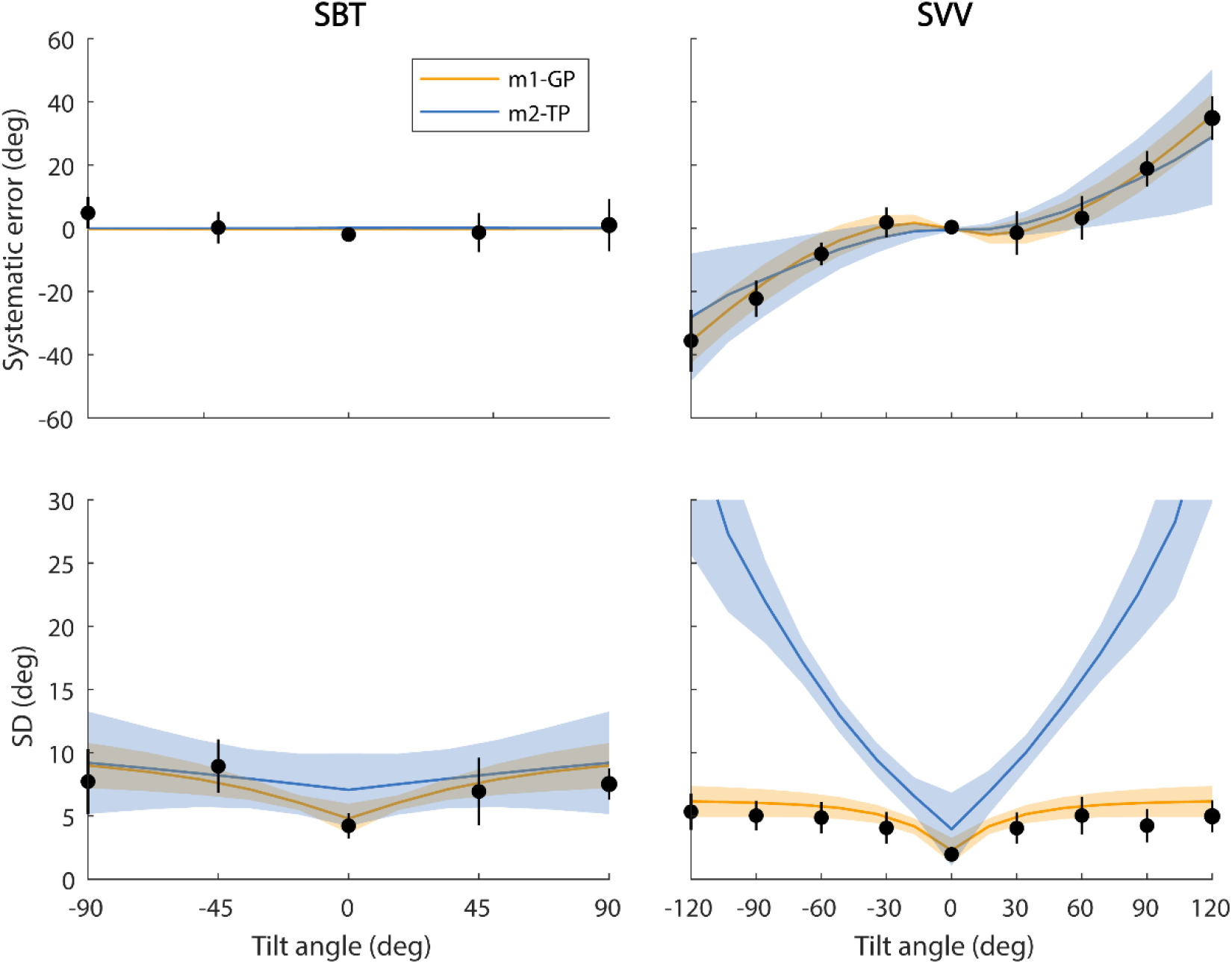
Predictions of the m1-GP (orange) and m2-TP models (*v* = 3.4, blue) of the SBT (left column) and SVV (right column), generated with the best-fitting parameter values per participant and then averaged across participants, plotted on top of the mean parameters from the psychometric fits (•). Shaded areas and error bars show one standard deviation above and below the participant mean.

**Figure 4.**
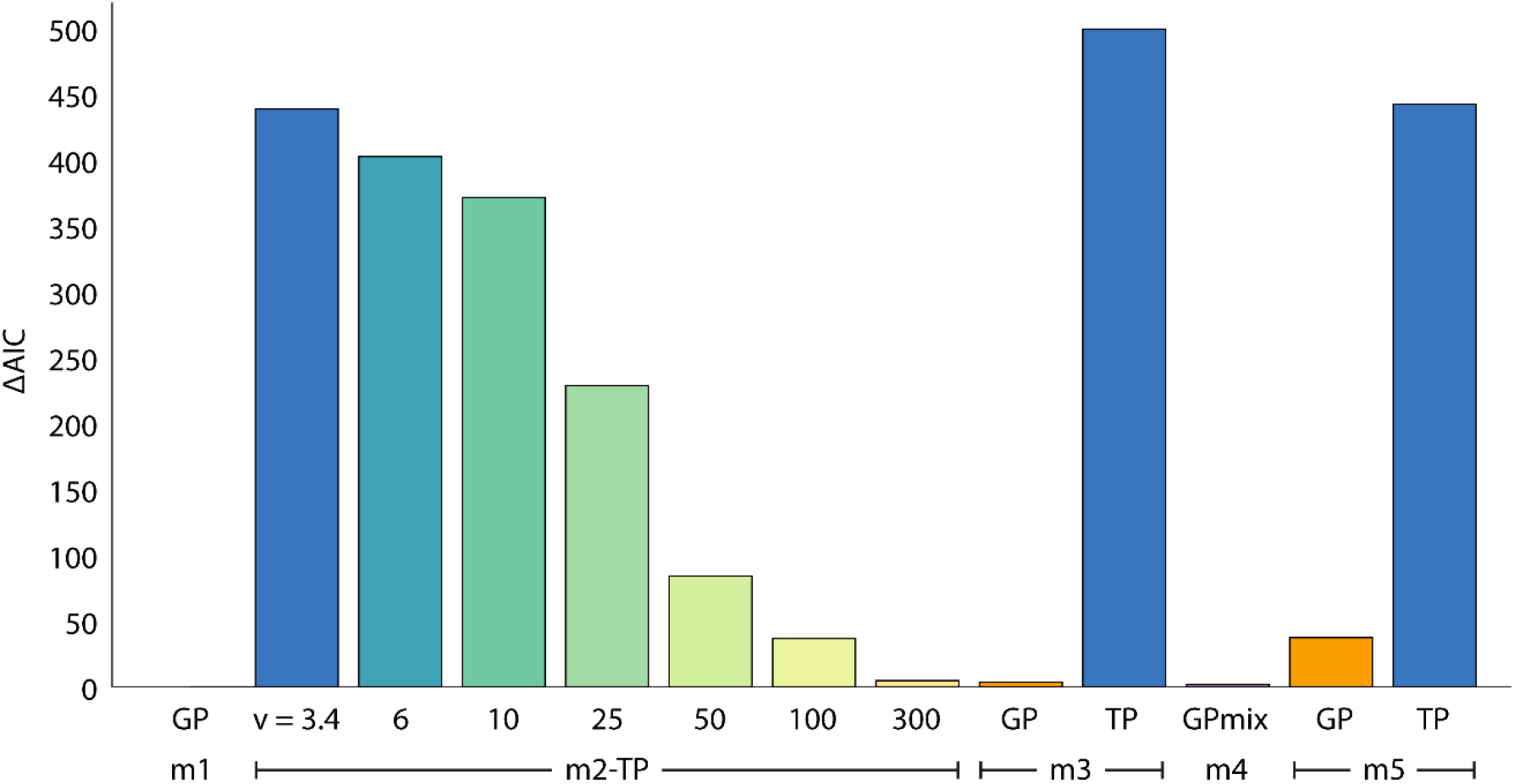
Comparison of the mean best AIC scores over participants of the different model variants, relative to the mean AIC score (439.2) of the Gaussian-prior model (m1-GP). A lower value indicates a better fit to the data. The different model variants are explained above (see Methods, Modeling, Model variants and their evaluation).

##### Model variant 1: m1-GP

This GP model is the numerical version of the original model by Clemens et al. (1) (from here on denoted as the m1-GP model). The original model contains 7 free parameters: *a_HS_, b_HS_, σ_BS_, σ_HB_, A_OCR_, λ* and *σ_HSP_* as the standard deviation of the Gaussian prior. Per participant, this model was fitted 100 times. For half of the fitting runs, random values within fixed bounds (*a_HS_*: [0, 0.5]°/°, *b_HS_, σ_BS_, σ_HB_, σ_HSP_*: [1e-05, 50]°, *A_OCR_*: [0, 30]°, *λ*: [0, 0.06]) were used as start values for the free parameters to maximize the possibility of convergence to the global minimum. The remaining runs were started with the fitted values from the Clemens et al. (1) study. Note, allowing a free parameter for the mean of the prior led to a fitted value close to 0. Therefore, this parameter was fixed at 0 during fitting.

##### Model variant 2: m2-TP

This TP model fitted the Clemens et al. (1) data set with the same free parameters but under the assumption of a *t-*location-scale prior (from here on denoted as the m2-TP model), which turned out to be the best description of the naturalistic head orientations in our data (see Figure 2B), corroborating previous literature (6). The shape parameter of the *t*-location-scale-prior distribution (*v_HSP_*) was fixed at the average parameter value of the best-fitting distribution on the naturalistic head orientation data (see Table S2 in the Supplemental Material). The fitting procedure was as for model 1. In further analyses, we also fitted this model using the shape parameter fixed at either 6, 10, 25, 50, 100 or 300 (resembling a Gaussian distribution), each fitted 50 times.

##### Model variant 3: m3-GP, m3-TP

In models 1 and 2 the standard deviation of the otolith noise depends linearly on the absolute head tilt. Instead, in model variant 3, we fitted a TP model without an imposed relationship between otolith noise and tilt angle, i.e., we allowed the standard deviation of the otolith noise to be a free parameter for each absolute tilt angle (from here on denoted as the m3-TP model). We used the fitted intercept and slope values of the Clemens et al. (1) study to compute a standard deviation for each tilt angle. These values were then used as the initial values for the free parameters in the fitting procedure, after which the parameters could take on any value within [1e-05, 50]°. We also repeated the fitting procedure with random start values for the free parameters. For comparison, this assumption was also tested for a Gaussian prior (the m3-GP model). Each version was fitted 100 times.

##### Model variant 4

This model variant involved a Gaussian-mixture distribution as head-in-space prior. A mixture of two Gaussians with the same mean but different SDs yields a distribution with heavier tails (to approximate the measured prior). We tested a prior distribution characterized by three parameters: the standard deviations of the two Gaussians, σ_HSP–1_ and σ_HSP–2_, and their mixing coefficient *c*, defined between 0 and 1, which weighs the two distributions. The means of both Gaussians were fixed at 0. The fitted values from Clemens et al. (1) were used as start values for the fitting procedure, where the fitted value for the standard deviation of the prior, σ_HSP_, served as start value for σ_HSP–1_. The start value for σ_HSP–2_ was 50°and the initial value of the mixing coefficient was valued either 0.25, 0.5, 0.75 or 1. The model was fitted 50 times for each of the different start values for the mixing coefficient.

##### Model variant 5: m5-GP, m5-TP

In the original model, the sensory measurements at a particular tilt angle are assumed to be unbiased on average but contaminated with independent Gaussian noise. This is referred to as a measurement distribution, i.e., the distribution of sensory tilt signals that is produced when the head is tilted at a specific angle. However, the brain must perform the inverse approach to find out which tilt angle has been responsible for the sensory signal that it receives. Hence it must compute the sensory likelihoods. If the measurement distribution of a sensory signal is Gaussian with a constant standard deviation, irrespective of tilt, the likelihoods will be Gaussian as well. For the otoliths, however, the standard deviation of the noise was assumed to be increasing with tilt angle, which formally results in a skewed otolith likelihood (see Girshick et al. (10) for a more detailed explanation). This nonlinear transformation was neglected in the original model – we assumed a symmetric otolith likelihood – but was put to test in variant 5. This model variant contained the same free parameters as m1-GP and m2-TP and was again fitted 100 times per prior form (from here on denoted as the m5-GP and m5-TP models).

## Results

We investigated the role of the prior’s form in a Bayesian model of spatial orientation, as assessed by the subjective visual vertical and subjective body tilt tasks at tilt angles between −120 to 120°. Data from these tasks were previously collected and extensively described and modelled using a Gaussian head-tilt prior in Clemens et al. (1). However, more recent work on head motion statistics reported that probability distributions of angular velocity and linear acceleration averaged across natural activities were not Gaussian, showing large positive excess kurtosis values (6). We first examined whether this observation can be generalized to head tilt distributions by measuring the head orientation statistics of human participants during typical everyday activities, and subsequently tested whether the model fit of the Clemens et al. (1) data can be improved by canceling the restriction to a Gaussian prior and allowing prior distributions more akin to these naturalistic head orientation distributions.

### Natural head orientation statistics

Figure 2A shows head orientation as a function of time for the various activities (i.e., walking, running, going up and down the stairs, sitting, and standing), separately for each participant. We found that the recorded head orientations varied across the activities. For example, over consecutive samples the changes in head tilt were smaller during sitting and standing than during the other activities. Furthermore, we found that probability distributions of head orientation pooled across activities were not Gaussian (Figure 2B) as quantified by large (between 5 and 142) kurtosis values across participants (Figure 2B insets). This indicates that head tilt distributions have longer tails and a higher peak than would be expected from normally distributed data. Across participants the ranges of the first three statistical moments of the head orientation data were *M* = −2.9° – 1.0°, *SD* = 6.3° – 10.3° and *S* = −2.1 – 3.2, indicating that the head tilt distribution centers on upright and shows no systematic skewness.

We next tested for each participant which of several probability distributions – the Gaussian, logistic, *t*-location-scale, extreme value and generalized extreme value distributions – best accounted for the measured head orientation statistics. We found that, pooled across activities, a *t*-location-scale distribution provided the best fit for five of the six participants, outperforming the Gaussian distribution in all cases (compare solid and dashed lines in Figure 2B). The relatively skewed data of participant 2 are described best by the extreme value distribution, followed by the *t*-location-scale distribution. We refer to Table S1 in the Supplemental Material for a comparison of the AIC scores of all fitted distributions. The parameters of the fitted *t*-location-scale distribution are consistent across participants (see Table S2): the location parameter is close to 0 (range: −1.9° – 1.1°), indicating that participants held their head on average upright, the scale parameter ranged between 3.9 and 6.8°, and the shape parameter was small (range: 2.2 – 4.3), corresponding to a characterization in terms of long tails. The results are consistent with previously reported distributions of head velocity and acceleration (6, 21).

### Bayesian modeling of spatial orientation

#### Model variants 1 and 2

We subsequently tested the assumption that these natural statistics of head orientation are used as a prior in spatial orientation, and can account for the observations in the SVV and SBT tasks reported by Clemens et al. (1). Within the structure of their Bayesian model of spatial orientation (Figure 1), we compared the predicted performance in the SBT and SVV tasks under the assumption of a Gaussian-head-tilt prior (m1-GP) and a *t*-location-scale prior (m2-TP, shape parameter fixed at 3.4, which was the average best-fitting shape parameter value on the naturalistic head orientation data (see Table S2)).

Figure 3 shows these predictions as the average (±SD) of the individual best fits superimposed on the mean data across participants. The prediction of the m1-GP model replicates well the closed-form model fit. In contrast, the m2-TP model provides a poor fit, both with regard to the observed bias in the SVV (the Aubert effect) and, more prominently, its variance. Also the variance of the SBT seems not well accounted for by this TP model.

To compare the quality of various model variants, we computed their AIC scores, averaged across participants. The baseline in this comparison is the mean AIC score of the m1-GP model, set to zero at the left in Figure 4. As shown, the m2-TP model (*v* = 3.4) performs substantially worse than the m1-GP model.

Table 1 illustrates the best-fit parameter values for each participant for the m1-GP and m2-TP (*v* = 3.4) models as well as the fit parameters reported in Clemens et al. (1) based on the original closed-form implementation. Comparing the m1-GP model with the closed-form implementation reveals similar fitted parameter values, confirming the numerical implementation of the Clemens et al. (1) model. The best-fitting m2-TP model yields large inter-participant variability for most of the parameters, suggesting that this TP variant does not capture the data very well.

**Table 1.** Best-fitting parameter values for the original implementation of the model as presented in Clemens et al. (1) (O), the numerical model with a Gaussian prior (m1-GP) and the numerical model with the *t*-location-scale prior (m2-TP, *v* = 3.4). The mean parameter values and their respective standard deviations are computed by averaging across participants. The fit parameters had the following lower and upper bounds: *a_HS_*: [0, 0.5]°/°, *b_HS_, σ_BS_, σ_HB_, σ_HSP_*: [1e−5, 50]°, *A_OCR_*: [0, 30]°, *λ*: [0, 0.06] (fitted lapse rates not presented).

| Parameter | $a_{HS}$ ($^{\circ}/^{\circ}$) | | | $b_{HS}$ ($^{\circ}$) | | | $\sigma_{BS}$ ($^{\circ}$) | | |
| --- | --- | --- | --- | --- | --- | --- | --- | --- | --- |
| Model version | O | m1-GP | m2-TP | O | m1-GP | m2-TP | O | m1-GP | m2-TP |
| Participant 1 | 0.23 | 0.23 | 0.23 | 1.2 | 1.4 | 5.5 | 12.3 | 12.4 | 8.2 |
| 2 | 0.12 | 0.12 | 0.24 | 1.2 | 1.2 | 0.8 | 8.4 | 8.3 | 9.8 |
| 3 | 0.20 | 0.21 | 0.32 | 1.1 | 1.1 | 0.5 | 6.7 | 6.4 | 3.1 |
| 4 | 0.07 | 0.07 | 0.30 | 3.9 | 3.9 | 30.7 | 12.6 | 12.6 | 9.9 |
| 5 | 0.11 | 0.11 | 0.49 | 3.3 | 3.4 | 22.7 | 15.0 | 15.1 | 9.9 |
| 6 | 0.23 | 0.23 | 0.18 | 3.0 | 3.2 | 3.1 | 8.0 | 8.2 | 6.1 |
| 7 | 0.20 | 0.20 | 0.18 | 3.2 | 3.4 | 2.5 | 12.7 | 12.6 | 40.8 |
| Mean ± | 0.16 ± | 0.17 ± | 0.28 ± | 2.4 ± | 2.5 ± | 9.4 ± | 10.8 ± | 10.8 ± | 12.5 ± |
| SD | 0.06 | 0.06 | 0.11 | 1.2 | 1.2 | 12.2 | 3.1 | 3.2 | 12.7 |

Table 1. Best-fitting parameter values for the original implementation of the model as presented in Clemens et al.
| Parameter | $\sigma_{HB}$ ($^{\circ}$) | | | $A_{OCR}$ ($^{\circ}$) | | | $\sigma_{HSP}$ ($^{\circ}$) | | |
| --- | --- | --- | --- | --- | --- | --- | --- | --- | --- |
| Model version | O | m1-GP | m2-TP | O | m1-GP | m2-TP | O | m1-GP | m2-TP |
| Participant 1 | 3.3 | 3.3 | 47.5 | 27.0 | 26.9 | 7.0 | 11.6 | 11.5 | 38.8 |
| 2 | 6.4 | 6.7 | 49.9 | 17.0 | 17.1 | 11.5 | 9.4 | 9.7 | 18.0 |
| 3 | 9.3 | 10.4 | 30.4 | 17.5 | 17.6 | 10.0 | 14.4 | 15.3 | 30.4 |
| 4 | 7.1 | 7.1 | 38.3 | 0.0 | 0.0 | 30 | 11.2 | 11.2 | 15.5 |
| 5 | 3.6 | 3.6 | 38.9 | 1.0 | 1.0 | 29.7 | 18.7 | 18.8 | 17.6 |
| 6 | 1.8 | 2.0 | 35.9 | 18.8 | 18.8 | 2.4 | 9.5 | 9.6 | 35.5 |
| 7 | 3.0 | 3.0 | 1.5 | 20.8 | 20.7 | 1.8 | 12.8 | 12.9 | 21.2 |
| Mean ± | 4.9 ± | 5.2 ± | 34.6 ± | 14.6 ± | 14.6 ± | 13.2 ± | 12.5 ± | 12.7 ± | 25.3 ± |
| SD | 2.7 | 3.0 | 16.1 | 10.2 | 10.1 | 11.9 | 3.2 | 3.3 | 9.5 |

Within the context of the m2-TP model, the decay rate of the prior distribution is captured by the shape parameter. The larger the value of this parameter, the closer the distribution approximates a Gaussian distribution. Figure 5 illustrates the predictions of the m2-TP model with the shape parameter fixed at 6, 10, 25, 50, 100, and 300 and averaged across participants, superimposed on the prediction of the m1-GP model. The model fit clearly improves with a larger shape parameter, which is confirmed by the corresponding ΔAIC scores in Figure 4, suggesting that the model better operates as the *t*-location-scale prior approximates a Gaussian. This is in stark contrast with the shape parameter values of 2.2 - 4.3 that we observed in the natural head orientations.

**Figure 5.**
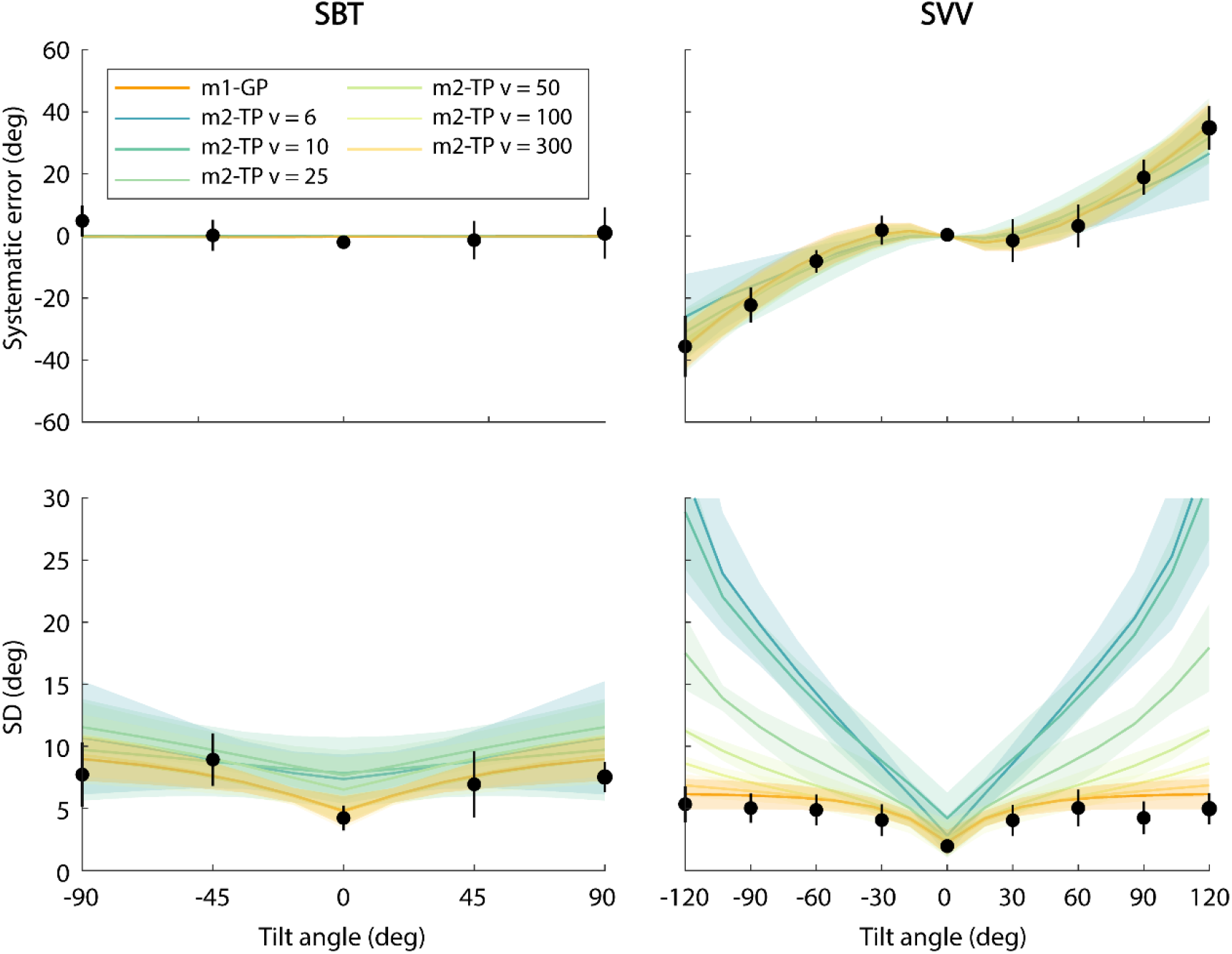
Predictions of the m2-TP model with different shape parameters (*v* = 6, 10, 25, 50, 100 or 300) and the m1-GP model, plotted on top of the mean parameters from the psychometric fits (•). Data are in the same format as in Figure 3.

#### Model variant 3

We next examined if releasing other model constraints can redeem the TP model. One constraint of the original model is that the standard deviation of the otolith noise depends linearly on the (absolute) tilt angle. In model variant 3 we lifted this constraint and fitted the GP and TP models with the standard deviation of the otolith noise a free parameter for each absolute tilt angle. With this additional flexibility, the m3-GP model still performed very well, but the m3-TP model did not improve, showing a higher ΔAIC score than without this flexibility (see Figure 4).

Figure 6 shows for both models the best-fitting values of otolith noise SD as a function of tilt angle, averaged across participants. For the GP model, there seems to be a linear relationship with absolute tilt angle, validating this constraint within the original model, while the TP model reveals a noisy, irregular pattern, resulting in a higher AIC score.

**Figure 6.**
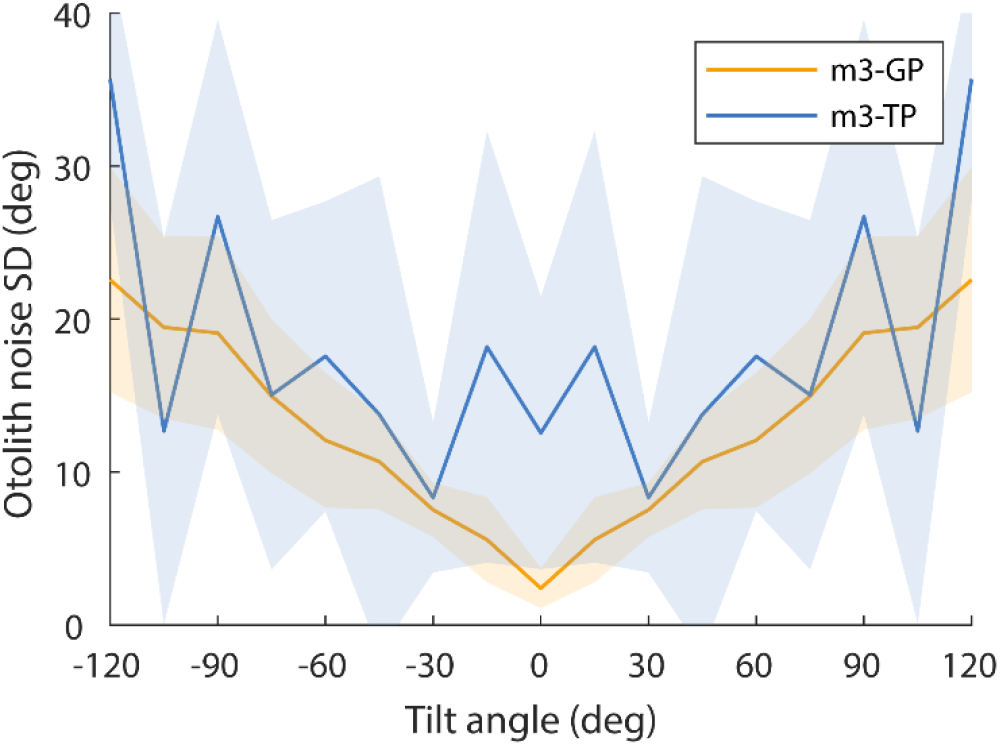
Best-fitting otolith noise values of the Gaussian-prior (orange) and *t*-location-scale-prior (blue) models as a function of tilt angle, averaged across participants. Shaded areas show one standard deviation above and below the participant mean.

#### Model variant 4

In model variant 4 we tested if a head-tilt prior that consists of a mixture of two Gaussians, which, compared to a *t*-location-scale distribution, can result in an alternative prior distribution with longer tails than a single Gaussian, fitted the data of Clemens et al. (1). The AIC score of the best-fitting Gaussian-mixture-prior model is comparable to the m1-GP model AIC score (but slightly higher, caused by the extra free parameters of the Gaussian-mixture prior (see Figure 4)). However, we found the mixing coefficients to be exactly 1, reducing the Gaussian-mixture distribution in fact to the single Gaussian distribution of the m1-GP model. Thus, the m1-GP model holds as the most parsimonious explanation of the data.

#### Model variant 5

Finally, we fitted a model variant that contains a skewed distribution for the otolith likelihood instead of the symmetric, Gaussian otolith likelihood in the original model. Again, the GP version of this model outperformed the same model with a *t*-location-scale prior. Furthermore, the m5-GP model performed worse than the original model in terms of AIC score (see Figure 4) and the m5-TP variant did not lead to an improved fit, showing a similar ΔAIC score as the m2-TP (*v* = 3.4) model.

## Discussion

The starting point of the present study is the Bayesian model of spatial orientation that we first proposed in 2011 (1). Specifically, a Gaussian-prior probability distribution of head roll was imposed to explain biases in the subjective vertical – known as the Aubert effect (see Figure 1). This prior probability distribution was regarded as a Bayesian observer’s assumption that the head is usually nearly upright (12, 28). Under the assumption that human observers are performing Bayesian inference for spatial orientation, we asked the question whether this form of the prior probability distribution is consistent with the natural statistics of head orientation, generated by human participants during everyday activities.

The answer is no. We found that the natural statistics of head orientation were poorly represented by Gaussian probability distributions but were characterized by long tails, as quantified by large kurtosis values. This observation extends observations by Carriot et al. (6) on head velocity and acceleration distributions. Their kurtosis values (>10) are similar to the range we found (between 5 and 14, with the exception of the much larger kurtosis value of participant 6). In both studies, these statistics are based on the combined distribution of all activities tested, even though the range of head orientations varied across the activities. Also separate analyses of the activities within subjects revealed distributions with excess kurtosis in nearly all cases (see Table S3), suggesting that the kurtosis does not originate from sampling from different Gaussian distributions. This does not seem a divergent finding, as Schwabe and Blanke (22) also reported deviations of normality of measured head pitch of human participants when they were standing, walking around, or moving as if they were playing tennis. Similar observations were made in visual and auditory modalities (29–32). Also in songbirds, the distribution of the sung pitches is observed to have long, non-Gaussian tails (33).

To model the naturalistic head orientation data, we fitted several probability distributions – the Gaussian, logistic, *t*-location-scale, extreme value and generalized extreme value distributions – to the head orientation data of each participant. A Gaussian distribution was never the best fitting function, but was always outperformed by a *t*-location-scale distribution. The *t*-location-scale distribution approaches the Gaussian distribution as the shape parameter tends to infinity, whereas smaller values of the shape parameter yield heavier tails. The latter is what we observed. The best-fitting shape parameter ranged between 2.2 and 4.3 across our participants.

We further performed a Bayesian modeling analysis using the *t*-location-scale distribution of head roll as the prior. To this end, the original closed-form Bayesian model by Clemens et al. (1) was turned into a numerical version. While the numerical model could equally well account for response bias and variance in the subjective visual vertical and subjective body tilt tasks under the assumption of a Gaussian prior as the original model (see Figure 3), it failed dramatically with a *t*-location-scale prior (see ΔAIC scores in Figure 4). Indeed, the larger we allowed the shape parameter of the *t*-location-scale prior to be, i.e., the better it approximated a Gaussian, the better the model accounted for SVV and SBT performance. Adding more flexibility to the model by releasing constraints on the otolith likelihood did not improve the model with the *t*-location-scale prior form. Fitting the model with a Gaussian-mixture prior (an alternative prior distribution allowing long tails) returned a single Gaussian distribution as the best account. Also extending the model by including the nonlinear transformation between the otolith measurement distribution and likelihood failed to improve the *t*-location-scale prior fit (see Figure 4).

Given that natural statistics of head orientation are best characterized by a non-Gaussian distribution, why is SVV and SBT performance so much better accounted for by a Bayesian observer assuming a Gaussian head-in-space prior? We can only speculate about the answers to this question. First, the statistics of natural head motion may simply not be incorporated as a prior in such perceptual computations. Carriot et al. (6) have shown that the statistics of signals experienced during active movements differed from those experienced during passive movement. For instance, their participants experienced greater translational accelerations and angular velocities during active motion than passive motion. Typically, during active exploitation, the system relies heavily on sensory feedback to control our body to remain within limits of stability and to prevent falling. A *t*-distribution prior could define a ‘zone of stability’ – a movement-relevant prior to control the deliberate exploration of plausible motor commands that keeps the body within the borders of postural stability (cf. Zhou et al. (33)).

A more theoretical explanation arises if one realizes that observing new evidence not always reduces uncertainty under Bayes’ rule. In other words, a posterior distribution does not necessarily have a lower variance than the prior or likelihood distributions it is based upon. It can be shown that this holds in the case when the prior and likelihoods are Gaussian, but not in all other cases (34). Figure 7 illustrates this point. The posterior that follows from a Gaussian likelihood and *t*-location-scale prior *can* have a larger variance than either prior or likelihood, consistent with the predictions in Figure 3, where the SD of the SVV posterior is much larger with a *t*-location-scale prior than with a Gaussian prior. This increase in width (or decrease in precision) occurs when the distance between the prior and likelihood means becomes large enough. Hence, a *t*-location-scale prior can lead to a negative information gain (34), which is a situation that the brain may want to prevent. Instead, the posterior yields a smaller variance if it follows from a Gaussian prior. Gaussian priors will reduce the variance of the posterior across all Gaussian sensory likelihoods, thus creating a positive information gain (irrespective of the distance between the prior and likelihood distribution). New sensory evidence will thus decrease the system’s uncertainty about the state it has adopted. In functional terms, for vertical perception, a Gaussian prior therefore amounts to a particular precision-accuracy trade-off across the tilt range: it suppresses uncertainty at the expense of a systematic bias at larger tilt angles (12). This cannot generally be achieved with a *t*-location-scale prior in the context of the structure of the model by Clemens et al. (1). We do not argue that no other model structures can be conceived that deal with this notion, but such conceptual analysis goes beyond the present investigation.

**Figure 7.**
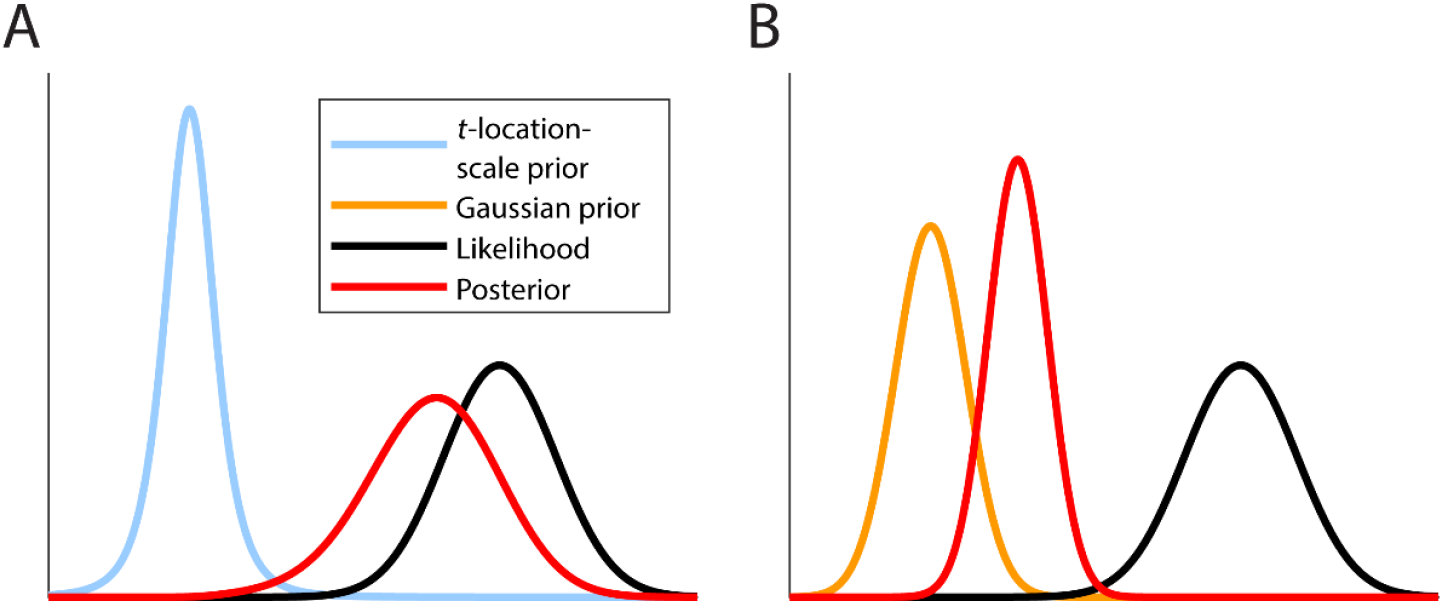
***A***: Integrating a *t*-location-scale prior (blue) which resembles the natural statistics of head orientations, with a Gaussian likelihood distribution (black) can lead to a posterior distribution (red) with a larger variance than the variances of the individual signals in the integration. ***B***: Instead, the integration of a Gaussian prior (orange) and Gaussian likelihood (black) always results in a Gaussian posterior (red) with a lower variance than the variances of the individual signals.

The subsequent question then is, how can the brain develop a Gaussian head-in-space prior, while the natural head motion statistics are best approximated by a *t*-location-scale distribution? As in any biological system, neural variability plays a role in vestibular processing and determines the neural code at central levels (35). Therefore, the signals at the level of the vestibular afferents will be noisier than the signals measured by the inertial measurement units, which will be close to the actual physical orientations. In other words, vestibular processing of head orientation signals is corrupted by additive (or multiplicative) noise (36). Based on the central limit theorem, this could, over an extended exposure to natural stimuli and daily-life tasks, convert the heavy-tailed distribution of measured head orientations into a more Gaussian distribution at the central level. If so, our results suggest that the brain stores this information and uses it as a prior in sensory processing for vertical perception.

At the neural level, vestibular afferents transmit data to the brain in trains of action potentials, and the brain needs to decode this information in terms of the head orientation, as well as other kinematic variables of head motion. It has been suggested that regular afferents transmit more information about changes in static head orientations than irregular afferents (35, 37). The likelihood distribution of the head’s kinematic state at the time of a spike of a given neuron differs from the prior distribution of states (38). Because a single spike transmits only a small amount of information, the observer’s uncertainty about the head’s kinematic state will be reduced (i.e., the variance of the posterior distribution will be smaller) if the prior distribution of states has a Gaussian form. More generally, it has been suggested that Bayesian computations with prior probabilities can rely on population vector decoding of neural populations with non-uniform preferred directions (10, 39).

As a final note, the considerations above assume the notion of a stable real-world prior, derived from the statistics of movements during natural activities over a long time. Stable priors have also been suggested for processing in other sensory modalities, e.g., predominance of horizontal and vertical orientations in natural scenes for visual orientation perception (5, 8, 10, 40). However, priors could also be more flexible, or context-dependent, and adapt over a short time scale, as for example has been shown in perceptual (41), motor (42) or language learning experiments (43). It remains to be tested how participants can develop a context- or task-dependent prior based on the orientations experienced during the experiment. The optimal strategy in this case is called dynamic or sequential Bayesian inference, which assumes conditionally independent measurements and Markovian dynamics. Under a recursive structure, it minimizes uncertainty in task outcome or state, given all measurements up to the present time or trial, by using the posterior distribution given all previous measurement as the prior distribution for inferring the posterior (33, 44–46). More specifically, a prior belief is computed by prediction, requiring a kinematic forward model, and then the posterior is updated by combining the likelihood with the prior (47, 48). It would be an interesting avenue for future work to embed the spatial orientation model of Clemens et al. (1) in this framework to find out which specific kinematic model could explain the dynamic SVV and SBT.

## Supplemental Material

**Table S1.**
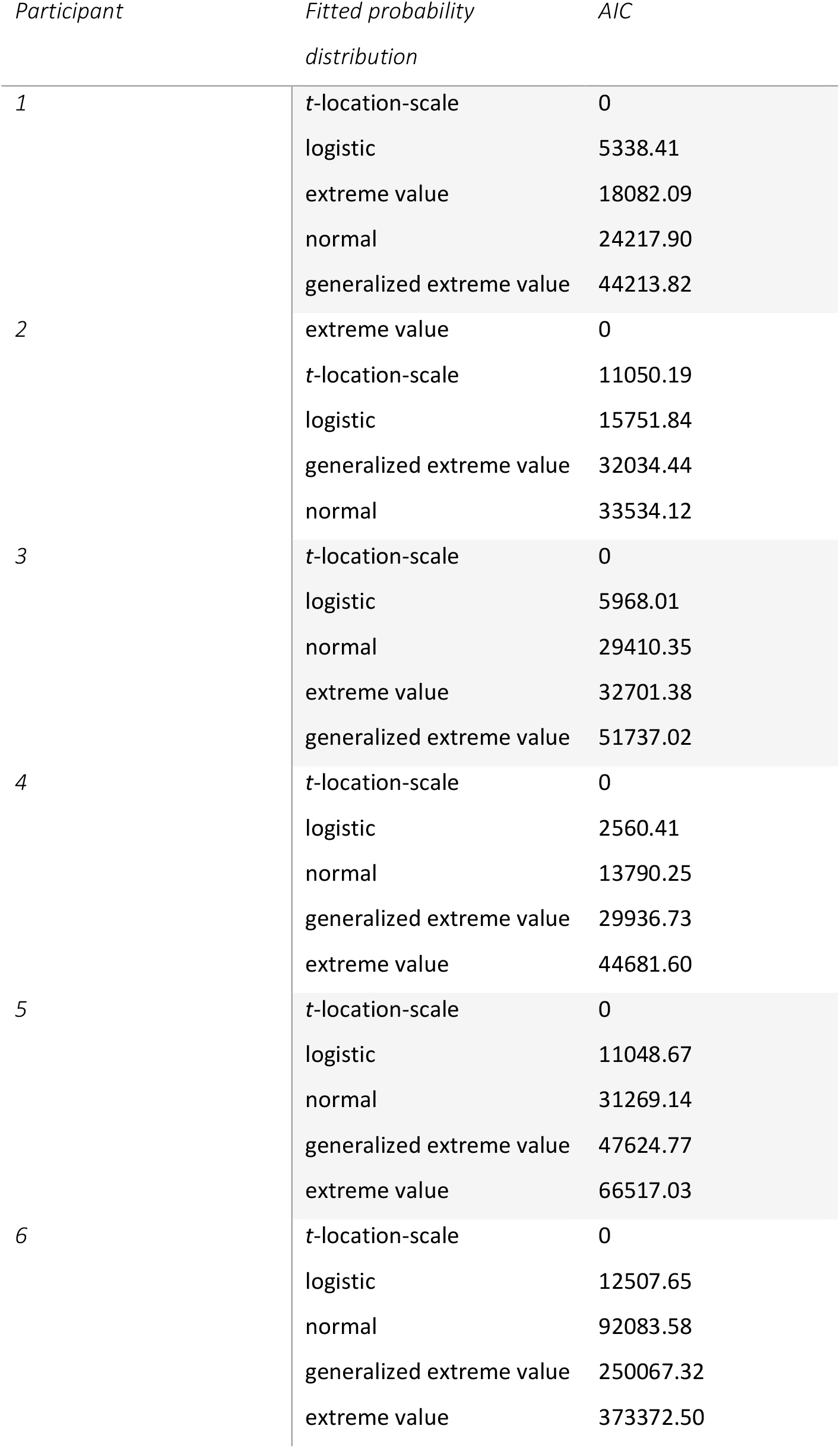
Best-fitting distributions on the head roll-tilt data pooled across activities, per participant, and their respective AIC scores, relative to best fitting distribution.

**Table S2.** Location, scale and (in the case of the *t*-location-scale distribution) shape parameter values of the *t*-location-scale and normal distributions that were fitted on the data pooled across activities for each participant. The bottom row shows the mean location, scale and shape parameters of the distributions, averaged across participants. The mean value of the shape parameter is used in the model fitting (see Methods, Modeling, Model variants and their evaluation, Model variant 2).

| *Participant* | *Probability distributions* | *Location parameter (deg)* | *Scale parameter (deg)* | *Shape parameter* |
| --- | --- | --- | --- | --- |
| 1 | t-location-scale | -1.91 | 6.77 | 3.28 |
|  | normal | -2.86 | 10.32 |  |
| 2 | t-location-scale | -0.79 | 5.57 | 3.33 |
|  | normal | -2.27 | 8.48 |  |
| 3 | t-location-scale | 0.95 | 4.81 | 3.55 |
|  | normal | 0.25 | 7.23 |  |
| 4 | t-location-scale | 1.11 | 4.39 | 3.52 |
|  | normal | 0.86 | 6.29 |  |
| 5 | t-location-scale | 1.05 | 3.93 | 2.23 |
|  | normal | 1.04 | 7.37 |  |
| 6 | t-location-scale | -0.11 | 4.91 | 4.28 |
|  | normal | 0.00 | 8.69 |  |
| Mean | t-location-scale | 0.05 | 5.06 | 3.36 |
|  | normal | -0.50 | 8.06 |  |

**Table S3.** Kurtosis values of the data of each activity of each participant.

| *Participant* | *Walking* | *Running* | *Standing* | *Sitting* | *Stairs* |
| --- | --- | --- | --- | --- | --- |
| 1 | 7.24 | 11.27 | 7.23 | 2.83 | 6.68 |
| 2 | 8.95 | 3.99 | 12.89 | 26.80 | 2.46 |
| 3 | 13.70 | 9.26 | 23.85 | 7.11 | 5.36 |
| 4 | 5.31 | 4.37 | 4.48 | 5.20 | 3.79 |
| 5 | 15.86 | 4.17 | 14.07 | 57.37 | 3.51 |
| 6 | 5.59 | 4.16 | 83.99 | 3.56 | 5.50 |

## Acknowledgments

We thank Dr. Antonella Pomante for designing and performing the naturalistic motion tracking experiment and Lucas Billen, MSc, for his contribution to the data analysis of the natural head orientation statistics.

## Author contributions

S.C.M.J.W., L.O.W., R.J.v.B., M.K. and W.P.M. conceived and designed research; S.C.M.J.W., R.J.v.B. and M.K. analyzed data; S.C.M.J.W., L.O.W., R.J.v.B., M.K. and W.P.M. interpreted results; S.C.M.J.W. prepared figures; S.C.M.J.W. drafted manuscript; S.C.M.J.W., L.O.W., R.J.v.B., M.K. and W.P.M. edited and revised manuscript; S.C.M.J.W., L.O.W., R.J.v.B., M.K. and W.P.M. approved final version of manuscript.

## Data availability statement

Upon publication, all data and code will be made publicly available via the data repository of the Donders Institute for Brain, Cognition and Behaviour.

